# Gradient-independent Wnt signaling instructs asymmetric neurite pruning in *C. elegans*

**DOI:** 10.1101/715912

**Authors:** Menghao Lu, Kota Mizumoto

## Abstract

During development, the nervous system undergoes refinement process by which neurons initially extend an excess number of neurites, the majority of which will be eliminated by the mechanism called neurite pruning. Some neurites undergo stereotyped and developmentally regulated pruning. However, the signaling cues that instruct stereotyped neurite pruning are yet to be fully elucidated. Here we show that Wnt morphogen instructs stereotyped neurite pruning for proper neurite projection patterning of the cholinergic motor neuron called PDB in *C. elegans*. In the loss of *lin-44/wnt* and *lin-17/frizzled* mutant animals, the PDB neurites often failed to prune and grew towards the *lin-44-*expressing cells. Surprisingly, membrane-tethered *lin-44* is sufficient to induce proper neurite pruning in PDB, suggesting that neurite pruning does not require Wnt gradient. LIN-17 and DSH-1/Dishevelled proteins were recruited to the pruning neurites in *lin-44-*dependent manners. Our results revealed the novel gradient-independent role of Wnt signaling in instructing neurite pruning.

## Introduction

Development of the nervous system is a highly dynamic process which involves neurogenesis, cell migration, neuronal polarization and outgrowth of neuronal processes. It is also well known that neurons initially extend an excess number of neurites most of which will later be eliminated by the mechanism called neurite pruning (Riccomagno and Kolodkin, 2015; Schuldiner and Yaron, 2015). Failure in neurite pruning results in excess arborization and neuronal connectivity, which could underlie various neurological conditions including macrocephaly (Jan and Jan, 2010). In vivo imaging techniques have allowed for the elucidation of genetic and molecular mechanisms that underlie neuronal pruning.

During neuronal refinement, neurites often compete with each other for their target cells in an activity-dependent manner, and neurites that are outcompeted are eliminated by pruning. For example, each muscle fiber in the mouse trapezius muscle is initially innervated by multiple motor neurons in early postnatal stage, while axons with weaker synaptic strength are pruned by the mechanism called axosome shedding (Bishop et al., 2004; Colman et al., 1997). The neurites also compete for trophic factors. Axons from rodent sympathetic neurons receive nerve growth factor (NGF) secreted from the target cells for their survival by inducing the brain-derived neurotrophic factor (BDNF), which in turn induces pruning of the outcompeted axons (Deppmann et al., 2008; Singh et al., 2008).

In addition to activity-dependent pruning, many neurites undergo stereotyped pruning during development (Schuldiner and Yaron, 2015). Several signaling molecules have been shown to play critical roles in this process. In mouse hippocampus, a chemorepellent Semaphorin 3F (Sema3F) is expressed in the distal infrapyramidal region and locally induces infrapyramidal tract (IFT) axon pruning via the PlexinA3-Neuropilin2 receptor complex (Bagri et al., 2003; Sahay et al., 2003). Neuropilin2 binds and activates β2-Chimaerin, a RacGEF, to inactivate Rac1 GTPase to induce IFT axon pruning (Riccomagno et al., 2012). Interestingly, loss of β2-Chimaerin specifically affects the pruning of the IFT axons but does not mimic the other phenotypes of the Sema3F or PlexinA2 such as axon guidance. Therefore, Sema-Plexin may utilize a specific downstream signaling cascade to control neurite pruning that differs from its roles in axon guidance and synapse formation.

Neurite pruning also plays a crucial role in sexually dimorphic neurocircuit formation. Expression of BDNF from the mammary mesenchyme is required for the innervation of the mammary gland sensory axons. In males but not in females, a truncated TrkB receptor isoform is expressed in the mammary mesenchyme which deprives BDNF availability for sensory axons thereby inducing male-specific axon pruning (Liu et al., 2012). Recent work revealed that Sema-Plexin signaling acts as a pruning cue for these sensory axons, as Sema6A and Sema3D expression in the mammary gland act induce sensory axon pruning through PlexinA4 (Sar Shalom et al., 2019).

Wnt is a secreted morphogen whose gradient distribution plays critical roles in various steps of animal development (Bartscherer and Boutros, 2008; Yang, 2012). Wnt signaling is also crucial in the nervous system development including neurogenesis, neuronal polarization, neurite outgrowth, axon guidance and synapse formation (He et al., 2018; Inestrosa and Varela-Nallar, 2015; Park and Shen, 2012). Nevertheless, its implication in neurite pruning is still limited. In *Caenorhabditis. elegans*, it has been shown that the transcription factor MBR-1/Mblk-1 is required for the pruning of the excess neurites of the AIM interneurons (Kage et al., 2005), whose function is antagonized by the trophic role of Wnt signaling (Hayashi et al., 2009). It has also been shown that Wnt signaling functions as a ‘pro-retraction’ cue in the Drosophila contralaterally-projecting serotonin-immunoreactive deuterocerebral interneurons (CSDns) (Singh et al., 2010). It is yet to be determined whether Wnt signaling plays instructive or permissive roles in neurite pruning.

In the present study, we describe a gradient-independent Wnt signaling in instructing stereotyped developmental neurite pruning of the PDB cholinergic motor neuron in *C. elegans.* From genetic analyses, we found that expression of LIN-44/Wnt in the tail hypodermal cells instructs pruning of the PDB neurites growing towards the tail hypodermal cells. Surprisingly, membrane-tethered LIN-44/Wnt is sufficient to induce PDB neurite pruning, suggesting that the gradient distribution of LIN-44 is not essential for neurite pruning. During this process, LIN-44/Wnt directs asymmetric localization of its receptor LIN-17/Frizzled and the intracellular signaling component DSH-1/Dishevelled to the posterior pruning neurites. Taken together, we discovered the novel gradient-independent Wnt signaling in instructing developmental neurite pruning.

## Results

### Asymmetric neurite pruning during PDB development

PDB is a post-embryonic cholinergic motor neuron derived from the P12 neuroectoblast cell (Sulston and Horvitz, 1977), and its cell body resides in the preanal ganglion. In hermaphrodites, PDB sends a single neurite posteriorly toward the tail tip region, where it makes sharp ‘V-shape’ turn to join the dorsal nerve cord to form *en passant* synapses onto dorsal body wall muscles (Figure 1A). While the function of PDB has not been characterized, a recent study combining mathematical modeling and a cell ablation experiment revealed the role of PDB in motor coordination (Walker et al., 2017). To investigate the PDB neurite development, we first generated a transgenic strain in which the PDB neurite and presynapses are labeled by a membrane-tethered GFPnovo2, a brighter GFP variant (Hendi and Mizumoto, 2018), and a synaptic vesicle gene, *rab-3*, fused with mCherry expressed under the *kal-1* promoter (Bülow et al., 2002) (see Materials and methods) (Figure 1A).

**Figure 1.**
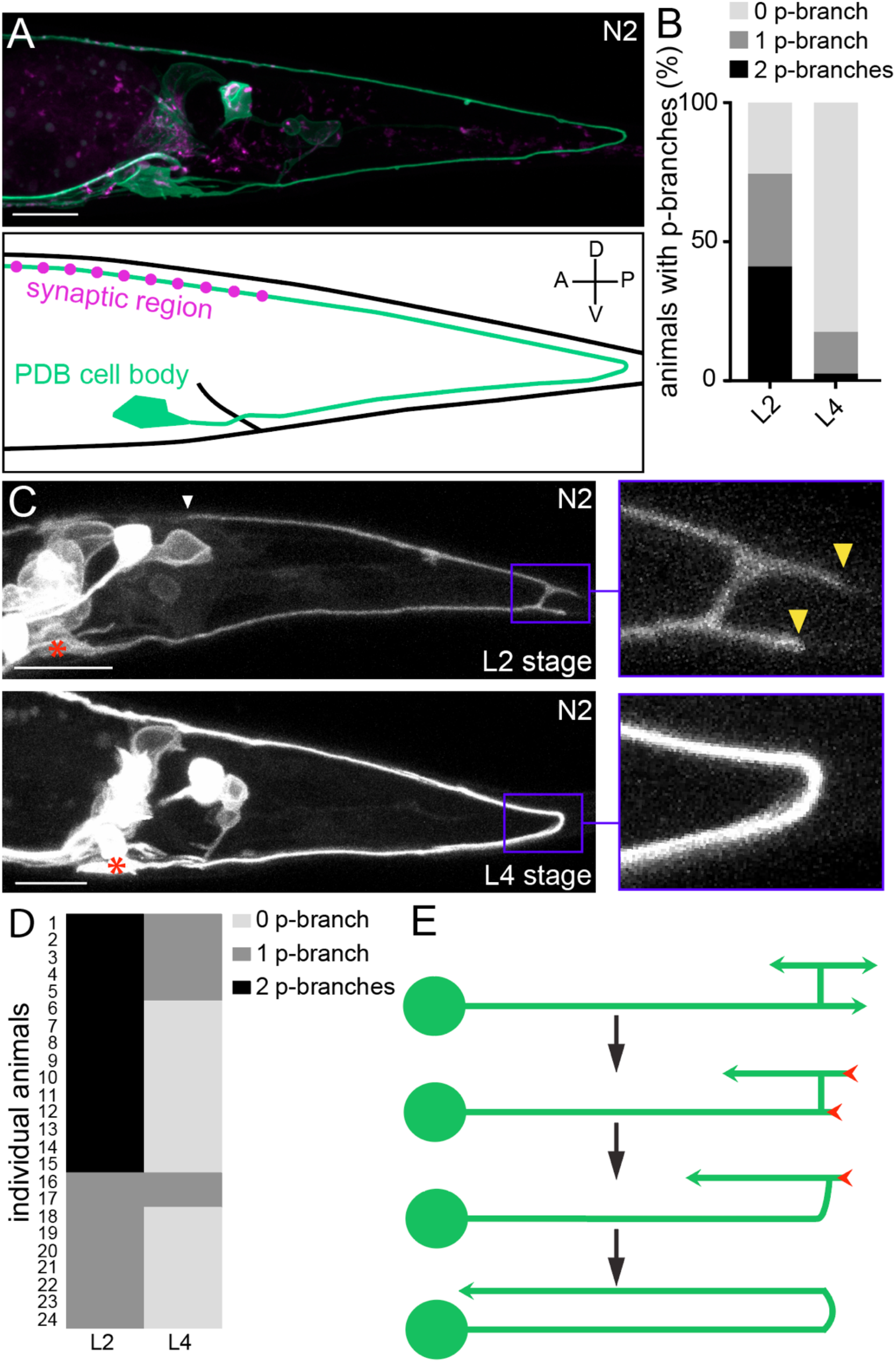
PDB neurites undergo stereotyped asymmetric pruning during development. (**A**) Structure of PDB labeled by *mizIs9.* The PDB process and presynaptic sites are labeled with GFPnovo2::CAAX (green) and mCherry::RAB-3 presynaptic vesicle marker (magenta), respectively. (**B**) Quantification of the number of posterior neurites (p-neurites) in semi-synchronized N2 populations at L2 and L4 stages respectively. (**C**) Representative images of the posterior neurite pruning event in a single animal at L2 and L4 stages. The regions of posterior neurites are magnified in the right panels. Asterisks represent PDB cell body. White and yellow arrowheads denote the end of anterior and posterior neurites, respectively. (**D**) quantification of the posterior neurite number of 24 animals at L2 and L4 stages. (**E**) A schematic of asymmetric neurite pruning during PDB development. Green and red arrowheads represent growing and pruning neurites. Scale bars: 10μm.

The PDB neuron extends its single neurite during late L1 and L2 stages. To understand how PDB neurite achieves V-shape turning, we first examined the structure of the developing PDB neurites using a population of semi-synchronized animals (Figure 1B). Interestingly, we observed that the PDB neuron has a unique transient structure at L2 stage: the ventral neurite sends a tiny commissure to the dorsal side of the worm where it bifurcates to extend one neurite posteriorly and one anteriorly (Figure 1C: **upper panels**). The PDB neuron therefore often has ‘H-shape’ structure with two posterior neurites (one dorsal neurite and one ventral neurite) at the early L2 stage (Figure 1B). In sharp contrast, the PDB neuron of the most L3/L4 animals contained no posterior neurite (Figures 1A and 1B). This observation suggests that the posterior neurites observed at the L2 stage are the transient structures most of which are pruned by the L4 stage. To confirm that the posterior neurites that are present at L2 stage are indeed pruned during development, we selected 24 L2 animals with at least one posterior neurite in PDB and re-examined its structure at L3/L4 stage (24 hours after the first examination) (Figures 1C and 1D). Among fifteen animals which had two neurites at the L2 stage, ten animals had no posterior neurite, and five animals had one neurite at L3/L4 stage. Among nine animals with one posterior neurite at the L2 stage, seven animals had no posterior neurite and two animals had one neurite at L3/L4 stage. In total, 82.1% (32/39) of the neurites we observed at the L2 stage were pruned within 24 hours.

Neurons could prune their neurites either by retracting or severing them (Schuldiner and Yaron, 2015). To distinguish which mechanism the PDB neuron utilizes to prune its posterior neurites, we conducted the time-lapse imaging during posterior neurite pruning and found that the posterior neurite became shorter over the development while the dorsal anterior neurite kept growing (Supplemental Figure 1). This indicates that the posterior neurites are pruned by retraction rather than the severing followed by degradation. To determine if PDB pruning is dependent on neuronal activity, we examined mutants of *unc-13*, which is required for synaptic vesicle fusion and neurotransmitter release (Richmond et al., 1999). We did not observe any structural defects in PDB such as ectopic posterior neurites in *unc-13* mutants, suggesting that the PDB neurite pruning is not dependent on the neuronal activity (Figure 2E). Taken together, our observation indicates that the V-shape turn of the PDB neurite is achieved by the asymmetric pruning of the posterior but not the anterior neurites at the turning point during PDB development (Figure 1E).

**Figure 2.**
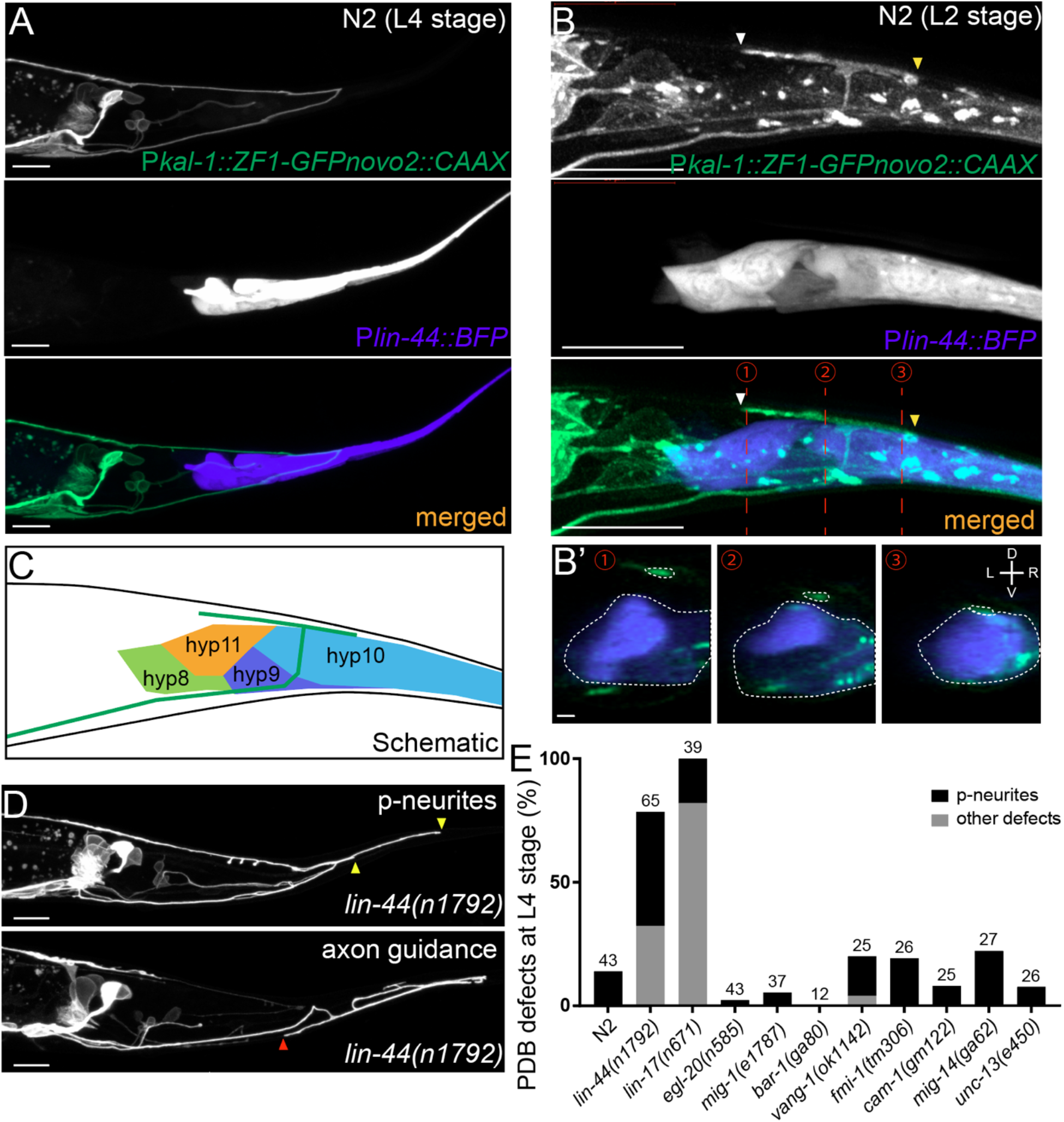
*lin-44/wnt* expressed adjacent to PDB posterior neurites is required for PDB development. (**A and B**) Representative images of PDB neurite labeled with *mizIs9* (top panels), *lin-44-*expressing cells labeled with P*lin-44::BFP* (middle panels) and merged images (bottom panels) at the L4 stage (**A**) and during posterior neurite pruning (L2 stage) (**B**). White and yellow arrowheads denote anterior and posterior neurites respectively. (**B’**) The transverse section of three positions of PDB neurites (indicated by red dotted lines in **B**) are reconstituted from the z-stack images shown in **B**. Dotted circles highlight PDB neurites and *lin-44-*expressing cells. (**C**) A schematic of (**B**). Green line represents PDB neurites. (**D**) Representative images of PDB with posterior neurites (top panel) and other defects (guidance defects: bottom panel) in *lin-44* mutants. Yellow arrowheads denote posterior neurites, and red arrowhead denotes anterior neurite failed to reach dorsal nerve cord. (**E**) Quantification of PDB defects at the L4 stage. Sample numbers are shown above each bar. Scale bars: 10μm.

### LIN-44/Wnt is required for the asymmetric neurite pruning in PDB

We next sought to identify the molecular cue that instructs pruning of the posterior neurites in PDB. Recent work has shown that the guidance and outgrowth of the PDB neurite are disrupted in the mutants of Wnt signaling and syndecan proteoglycans (Saied-Santiago et al., 2017). Wnt is a family of highly conserved secreted glycoproteins that play pivotal roles in animal development (Nusse, 2005). Wnt acts through receptors including Frizzled, LRP5/6 and receptor *tyrosine* kinase (Ryk). Upon Wnt binding, these receptors execute distinct downstream cascades via the multi-domain protein, Dishevelled (Sawa and Korswagen, 2013; van Amerongen and Nusse, 2009).

Previous works have shown that *lin-44/wnt* is expressed in the hypodermal cells at the tail tip region (Figure 2A), where it inhibits axon outgrowth and synapse formation, as well as controls neuronal polarity and asymmetric cell division (Herman et al., 1995; Hilliard and Bargmann, 2006; Klassen and Shen, 2007; Maro et al., 2009). We therefore examined the relative position of the PDB neurites and the *lin-44-*expressing cells labeled with the blue fluorescent protein (BFP) expressed under the *lin-44* promoter during development. Strikingly, at the L2 stage when PDB posterior neurites undergo pruning, we found that the pruning posterior neurite was located in close proximity to the most posterior hypodermal cell, hyp10 (Figures 2B and 2C). Interestingly, we noticed that hyp10 often showed the brightest BFP expression among the *lin-44-*expressing cells at the L2 stage. The position of the PDB posterior neurites and hyp10 which expressed a high level of *lin-44* is well in line with our prediction that *lin-44/wnt* is a strong candidate as a pruning cue. We then examined whether the pruning of the posterior neurites is affected in the *lin-44* null mutant background. At L4 stage, approximately 32.3% (21/65) of the *lin-44(n1792)* null animals showed cell fate determination defects (no PDB), neuronal polarity defects (PDB neurite extended anteriorly) and axon guidance defects (Figure 2D, **bottom panel and Data not shown**). We excluded them from our quantification because these phenotypes prevented us from examining the neurite pruning defects. Among the *lin-44* mutant animals with no obvious cell fate, neuronal polarity or guidance defects, 68.2% (30/44) of them showed ectopic posterior neurites in PDB at the L4 stage, suggesting the defective neurite pruning during PDB development (Figures 2D and 2E). To test if the pruning of the posterior neurites is compromised in the *lin-44* mutants, we compared the PDB structure of the same animal at two developmental time points, second larval (L2) and last larval (L4) stages (Figures 3A and 3D). As expected, we indeed observed that only 22.9% (8/35) of the posterior neurites observed at the L2 stage animals are pruned by L3/L4 stage in the *lin-44* mutants, which is significantly lower than those observed in wildtype (82.1%) (Figures 3C and 3D). This result strongly suggests that *lin-44/wnt* is required for the pruning of the posterior neurites during PDB development.

**Figure 3.**
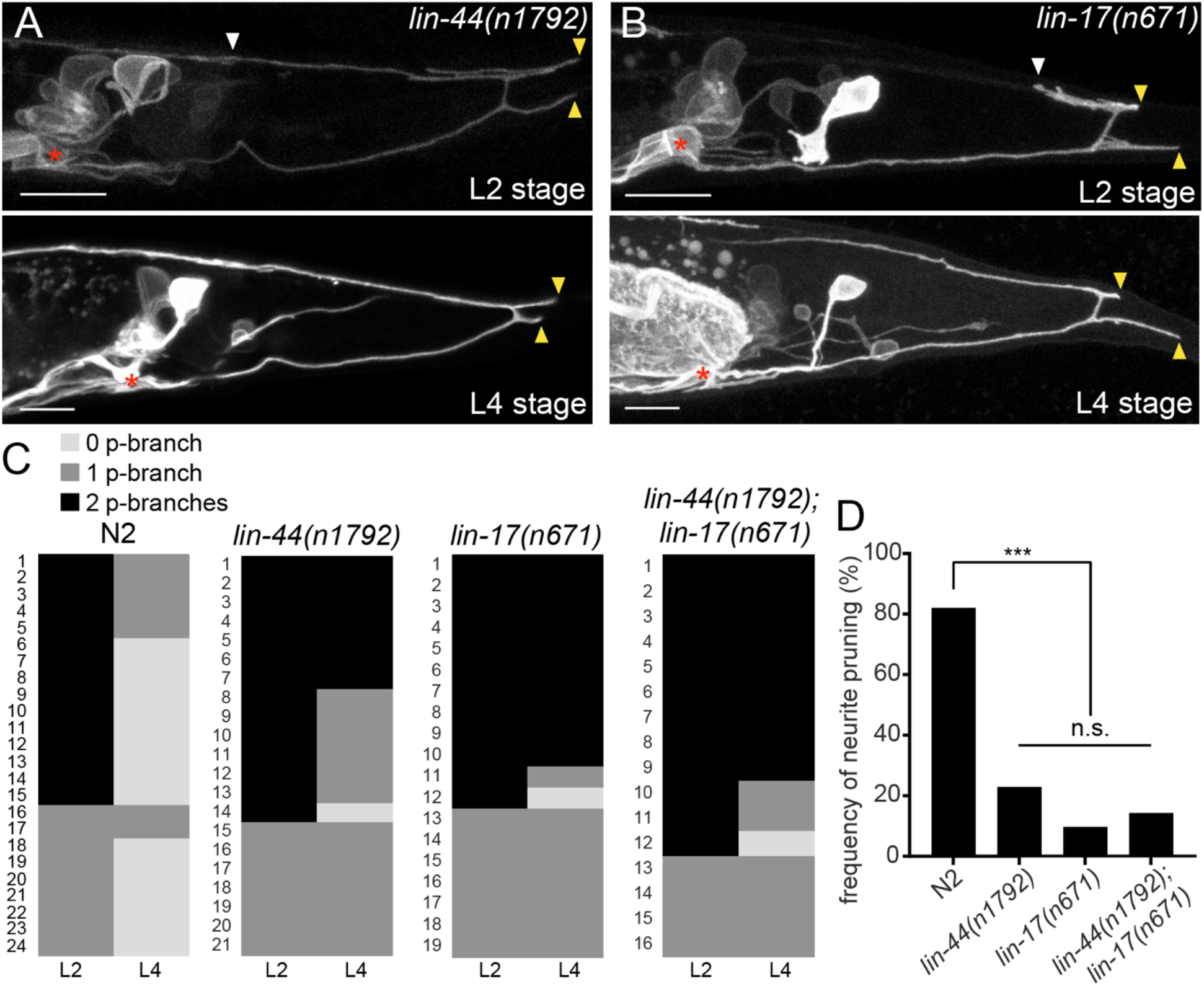
*lin-44/wnt* and *lin-17/fz* are required for the posterior neurite pruning. (**A** and **B**) Representative images of single animals of *lin-44(n1792)* (**A**) and *lin-17(n671)* (**B**) mutants at L2 (top panels) and L4 (bottom panels) stages. Asterisks represent PDB cell body. White and yellow arrowheads denote anterior and posterior neurites respectively. (**D**) Quantification of the posterior neurite number of 24 animals at L2 and L4 stages. Note that quantification of N2 is from Figure 1. (**D**) Quantification of the posterior neurite pruning frequency. *** p < 0.001; n.s., not significant (Chi-square with Yates’ correction). Scale bars: 10μm.

### *lin-17/fz* acts cell-autonomously in PDB to induce neurite pruning

LIN-44/Wnt utilizes LIN-17/Frizzled(Fz) as a primary receptor in many contexts (Hilliard and Bargmann, 2006; Klassen and Shen, 2007; Sawa et al., 1996). We then tested if *lin-17/fz* also plays a role in neurite pruning in PDB. While *lin-17(n671)* null mutants showed significantly more severe cell fate, polarity or guidance defects than *lin-44/wnt* mutants (Figure 2D), we observed similar posterior neurite pruning defects as *lin-44* mutants: only 9.7% of the posterior neurites (3/31) were pruned 24 hours after L2 stage (Figures 3B-3D). The pruning defect in the *lin-44; lin-17* double mutants was not significantly different from each single mutant (14.3% of the posterior neurites were pruned), suggesting that *lin-44* and *lin-17* act in the same genetic pathway (Figures 3C and 3D). This result is consistent with the idea that *lin-17/fz* is the main receptor for *lin-44/wnt* in PDB neurite pruning. We also conducted tissue-specific rescue experiments to test if *lin-17* functions cell-autonomously in PDB. Expression of *lin-17* cDNA either from the pan-neuronal promoter (P*rgef-1*) or the PDB promoter (P*kal-1*) significantly rescued the pruning defects of *lin-17* mutants, suggesting that *lin-17* acts cell-autonomously in PDB (Supplemental Figure 2). These results indicate that LIN-17/Fz is a receptor that functions in PDB to receive LIN-44/Wnt signal to induce neurite pruning.

### LIN-44/Wnt but not EGL-20/Wnt instructs PDB neurite pruning

Since *lin-44/wnt* is expressed posteriorly to the PDB neurites, it is possible that LIN-44 acts as an instructive cue for PDB neurite pruning. Alternatively, it might also act as a permissive cue for PDB neurite pruning. In order to distinguish these two possibilities, we conducted the rescue experiment by expressing LIN-44 ectopically using the promoter of another *wnt* gene, *egl-20*. *egl-20/wnt* is expressed in the cells around preanal ganglions (Whangbo and Kenyon, 1999). If LIN-44/Wnt functions as an instructive cue, its expression from anterior cells using the *egl-20* promoter would not rescue the PDB pruning defect in *lin-44* null mutants. If it acts as a permissive cue, we expect to see the rescue of the PDB neurite pruning defects no matter where *lin-44* is expressed. The *lin-44* mutants with an extrachromosomal array Ex[P*egl-20::lin-44*] did not rescue pruning defects of *lin-44* null mutants (31.8%, n = 22) (Figures 4A and 4B). On the other hand, the transgene expressing *lin-44* from the *lin-44* promoter (Ex[P*lin-44::lin-44*]) completely rescued the pruning defects (80%, n=20) to the wildtype level (Figures 4A and 4B). This result suggests that *lin-44/wnt* is an instructive cue for the posterior neurite pruning in PDB.

**Figure 4.**
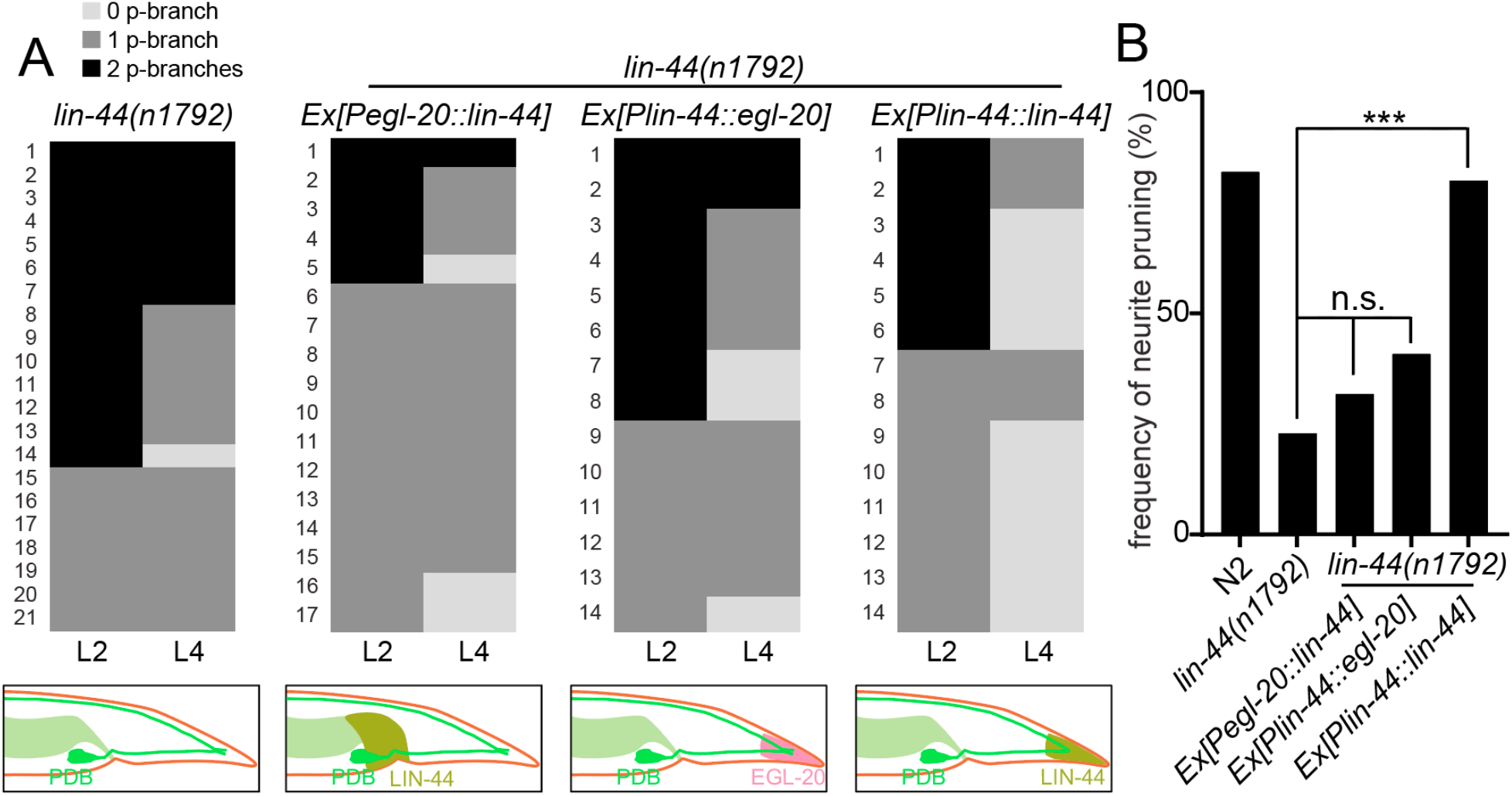
*lin-44* but not *egl-20* instructs neurite pruning in PDB. (**A**) Quantification of the posterior neurite numbers of individual animals at L2 and L4 stages in *lin-44* mutants and *lin-44* mutants with rescuing transgenes. Note that the quantification of *lin-44* mutants is from Figure 3. Bottom panels are schematics showing the expression domain of *lin-44* (green) and *egl-20* (magenta) in each genotype. (**B**) Quantification of posterior neurite pruning ratio. *** p < 0.001; n.s., not significant (Chi-square with Yates’ correction).

*egl-20/wnt* often acts cooperatively and redundantly with *lin-44/wnt* in various developmental events (Mizumoto and Shen, 2013; Yamamoto et al., 2011). We therefore tested if *egl-20/wnt* can replace the function of *lin-44* in PDB neurite pruning. Interestingly, expression of *egl-20* under the *lin-44* promoter did not rescue the PDB neurite pruning defects of the *lin-44* null mutant animals, suggesting the functional divergence between *lin-44* and *egl-20* in neurite pruning (Figures 4A and 4B). Consistently, PDB structure is largely unaffected in the mutants of *egl-20(n585)* and its receptors *mig-1/fz* and *cam-1/Ryk* (Figure 2E **and data not shown**) (Eisenmann, 2005; Mizumoto and Shen, 2013). Taken together, we conclude that LIN-44 is a unique instructive cue for PDB neurite pruning.

### Membrane-tethered LIN-44 is sufficient to instruct PDB neurite pruning

Genetic experiments such as ectopic expression or overexpression of Wnts, and direct visualization of the Wnt protein has revealed the critical roles of gradient distribution of LIN-44 and EGL-20 in *C. elegans* (Coudreuse, 2006; Klassen and Shen, 2007; Maro et al., 2009; Mizumoto and Shen, 2013; Pani and Goldstein, 2018). While data from the above experiments are consistent with the idea that *lin-44/wnt* instructs asymmetric neurite pruning in PDB, several observations cannot be explained by the instructive role of LIN-44 as a gradient. First, we did not observe ectopic pruning of the anterior neurite when *lin-44/wnt* was expressed anteriorly from the *egl-20* promoter (data not shown), suggesting that anterior expression of *lin-44* is not sufficient to induce anterior neurite pruning in PDB. Second, during PDB development, both growing anterior and pruning posterior neurites are located close to the *lin-44-*expressing hyp10 cell (Figure 2B). If *lin-44/wnt* functions as a gradient pruning cue, how can the anterior neurite escape from the pruning signal? One possible explanation is that PDB neurites require high and locally concentrated LIN-44/Wnt to induce pruning rather than responding to the gradient distribution of LIN-44. In this case, the cells expressing *Pegl-20::lin-44* are too far away from the anterior neurite of PDB. Furthermore, a close examination of the spatial relationship between PDB neurites and the *lin-44*-expressing cells suggests that the GFP signal from the PDB posterior neurite tip overlaps with the BFP signal from hyp10 while there is a gap between the anterior neurite tip and hyp10 (Figure 2B). This raises one possibility that PDB neurites need to be in very close proximity, if not in contact, with the LIN-44-expressing cells to activate the pruning mechanism. To test this hypothesis, we took advantage of the membrane-tethered Wnt originally developed in *Drosophila* (Zecca et al., 1996). In *Drosophila*, the fusion construct of the type-II transmembrane protein Neurotactin (Nrt) and Wingless/Wnt (Nrt-Wg) is used to characterize the gradient-independent role of Wingless/Wnt (Alexandre et al., 2014; Baena-Lopez et al., 2009). Importantly, the mutant fly in which endogenous *Wg* was replaced with *Nrt-Wg* was viable with no major morphological defects (Alexandre et al., 2014). They also showed that the Nrt-Wg largely abrogated the gradient distribution of Wingless. These observations suggest that the gradient distribution of Wnt is not essential for normal development. We generated the mutant animal with membrane-tethered *lin-44* by inserting the codon-optimized Neurotactin (Nrt) and BFP into the endogenous *lin-44* locus by CRISPR/Cas9 (*miz56*[*nrt-bfp-lin-44*]) (Figure 5A). The BFP signal was observed on the membrane of the tail hypodermal cells including hyp10, suggesting that NRT-BFP-LIN-44 fusion proteins are localized on the surface of the *lin-44-*expressing cells (Figure 5B). The brighter BFP signal was observed at the interface between the hypodermal cells, likely because the signal comes from both hypodermal cells. Interestingly, we observed significant proportions of the *nrt-bfp-lin-44* animals have normal PDB structures at the L4 stage (**data not shown**). Specifically, unlike *lin-44* null mutants, we observed normal pruning of the posterior neurites in *nrt-bfp-lin-44* animals (65.6%, n=32) comparable to wildtype (Figures 5C-5E). This indicates that the membrane-tethered LIN-44 is sufficient to induce normal neurite pruning in PDB.

**Figure 5.**
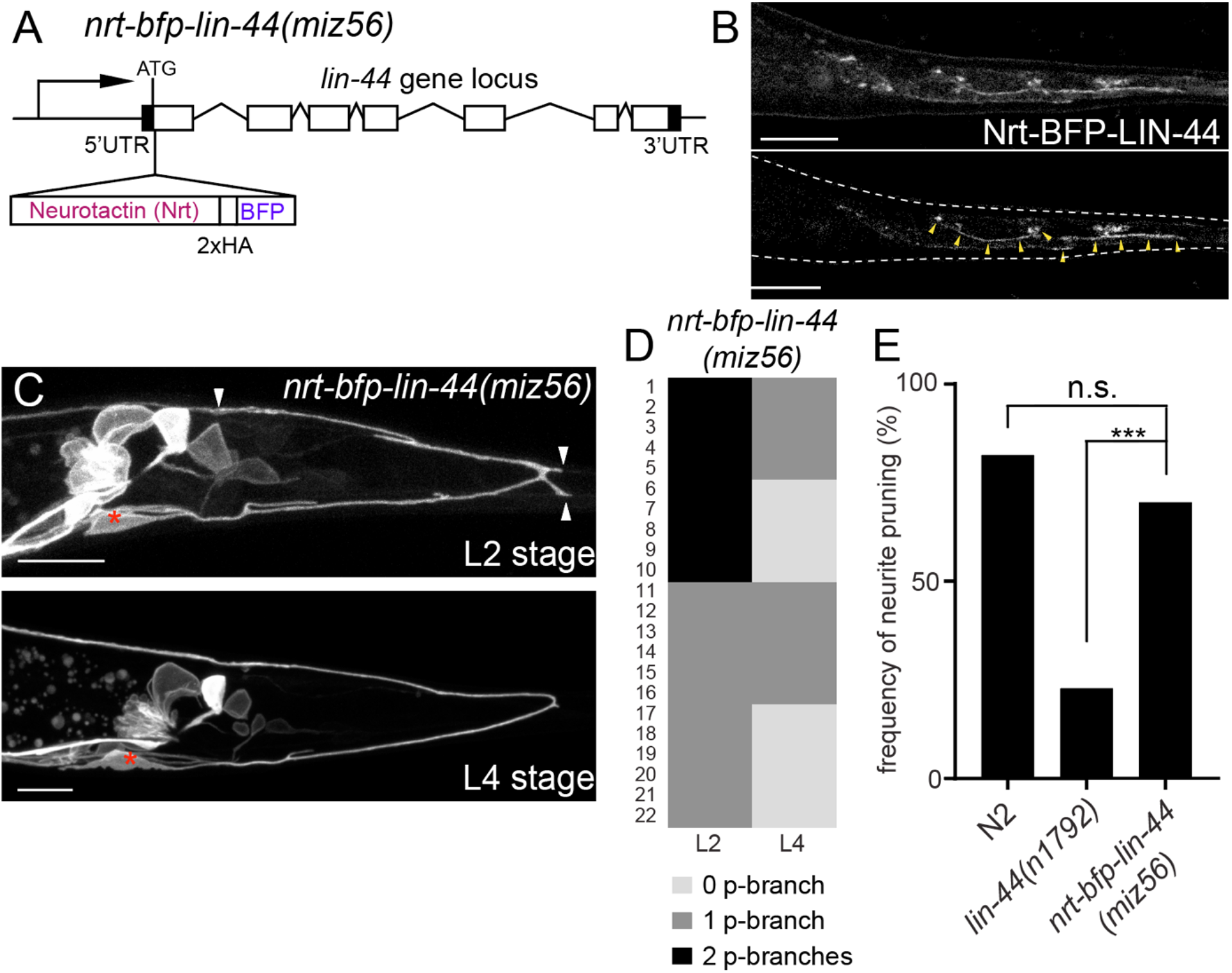
Membrane-tethered LIN-44 is sufficient to induce posterior neurite pruning in PDB. (**A**) A genomic structure of *lin-44* locus in *nrt-bfp-lin-44(miz56)* mutants. (**B**) Subcellular localization of Nrt-BFP-LIN-44 in the adult animal. Maximum projection (top panel) and single plane (bottom panel) from z-stack images. Arrowheads denote Nrt-BFP-LIN-44 signal on the membrane. (**C**) Representative images showing the pruning activity in *nrt-bfp-lin-44(miz56)*. (**D**) Quantification of the posterior neurite numbers of 22 individual animals at L2 and L4 stages in *nrt-bfp-lin-44(miz56)* mutants. (**E**) Quantification of posterior neurite pruning frequency. *** p < 0.001; n.s., not significant (Chi-square with Yates’ correction). Scale bars: 10μm.

In addition to the traditional model of Wnt gradient formation by simple secretion and chemical diffusion, recent works have started to reveal the alternative modes of Wnt transport via cytonemes and extracellular vesicles (EVs) (Gross et al., 2012; Routledge and Scholpp, 2019; Saha et al., 2016; Stanganello et al., 2015). It is therefore possible that the membrane-tethered Wnt is still capable of making a gradient and reach the target cells from distant via these structures. To test if NRT-BFP-LIN-44 abolished the gradient distribution of LIN-44, we examined the phenotypes of two motor neurons (DA9 and DD6) that are known to be regulated by the LIN-44/Wnt gradient signal. The DA9 neuron is a cholinergic motor neuron whose cell body resides in the preanal ganglion. DA9 axon initially extends posteriorly along the ventral nerve cord and through commissure join the dorsal nerve cord where it then extends anteriorly to form *en passant* synapses onto dorsal body wall muscles. Previous works have shown that LIN-44/Wnt gradient from the tail hypodermal cells inhibits synapse formation in the posterior axonal region of DA9, creating the synapse-free (or asynaptic) axonal domain (Figures 6A and 6D) (Klassen and Shen, 2007). In the *lin-44* null mutants, the ectopic synapse formation occurs in the dorsal posterior axonal domain of DA9 (Figures 6B and 6D). LIN-17::mCherry is localized within the asynaptic posterior axonal domain in a *lin-44-*dependent manner (Figures 6A and 6B, **middle panels**). Overexpression of *lin-44* from the *egl-20* promoter creates larger asynaptic domain by displacing DA9 synapses more anteriorly, suggesting the LIN-44 gradient determines the length of the asynaptic domain (Klassen and Shen, 2007). Similarly, the termination of the axonal outgrowth of the DD6 GABAergic motor neuron in the dorsal nerve cord is dependent on the LIN-44 gradient (Maro et al., 2009). In wildtype, the DD6 axon terminates before it reaches the rectum region (Figures 6E and 6F). In *lin-44* null mutants, it overextends beyond the rectum (Figures 6E and 6F). Overexpression of *lin-44* from the *lin-44* or *egl-20* promoters caused premature termination of the DD6 axon outgrowth (Maro et al., 2009). These two *lin-44*-dependent phenomena were severely disrupted in *nrt-bfp-lin-44* animals and were indistinguishable from those in *lin-44* null mutants (Figures 6C-6F). These results strongly suggest that membrane-tethered LIN-44 severely compromised the gradient-dependent function of LIN-44. These results are consistent with the idea that the pruning of the PDB posterior neurites does not depend on the LIN-44 gradient but on the local and high concentration of LIN-44 directly exposed by the *lin-44*-expressing cells.

**Figure 6.**
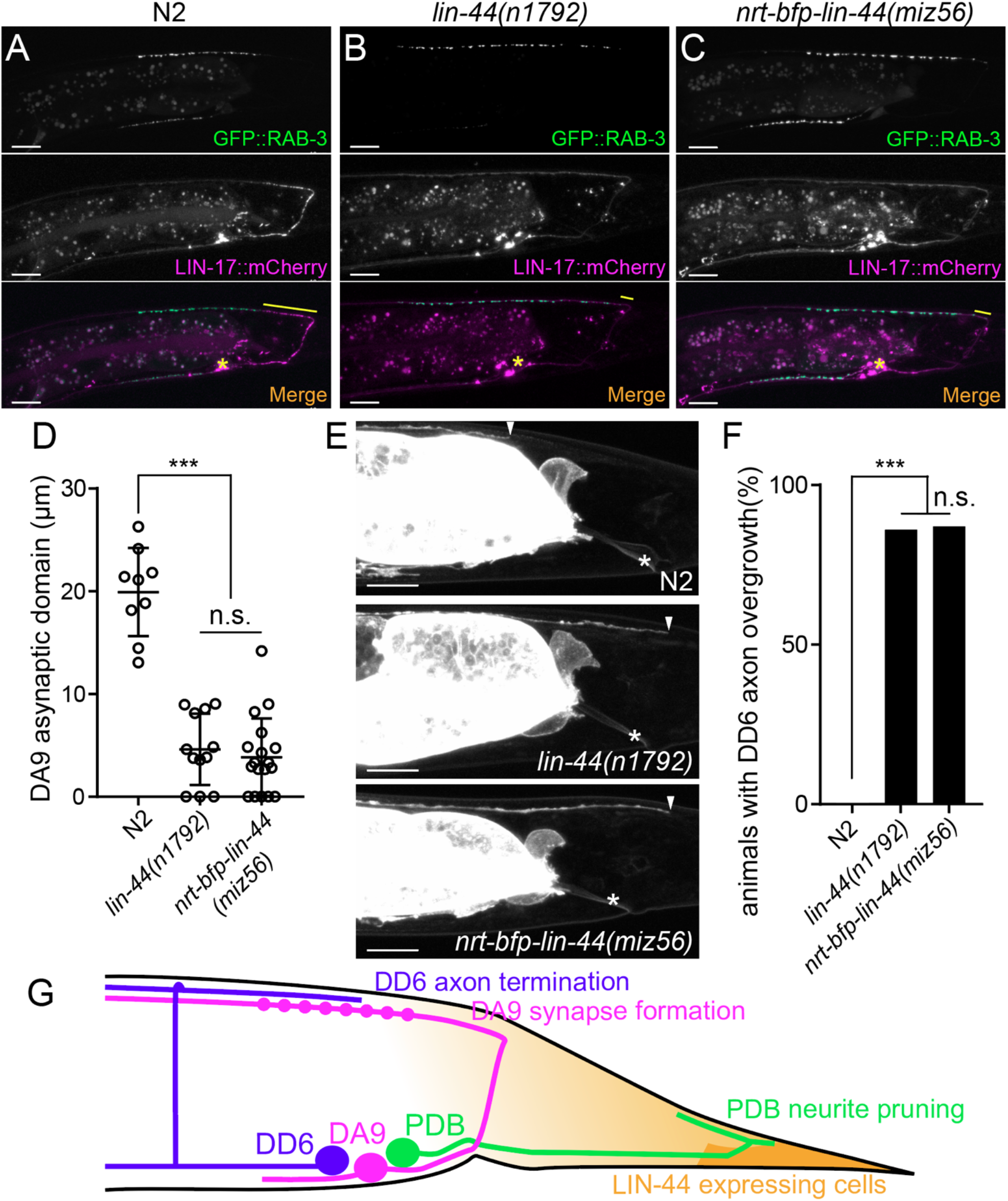
Membrane-tethered LIN-44 cannot function as a gradient signal. (**A-C**) Representative images of DA9 presynaptic specializations labeled with GFP::RAB-3 (top panels), LIN-17::mCherry localization (middle panels) and merged images (bottom panels) in N2 (**A**), *lin-44(n1792)* (**B**) and *nrt-bfp-lin-44(miz56)* (**C**) animals. Asterisks denote DA9 cell body, and yellow lines represent the posterior asynaptic domain of the DA9 dorsal axon. (**D**) Quantification of the DA9 asynaptic domain length. Each dot represents an individual animal. Error bars indicate mean ± SEM. ***p<0.001; n.s., not significant (one-way ANOVA). (**E**) Representative image of DD6 posterior axon in N2 (top panel), *lin-44(n1792)* (middle panel) and *nrt-bfp-lin-44(miz56)* (bottom panel). Asterisks denote the position of the rectum; arrowheads denote the end of DD6 axon. (**F**) Quantification of DD6 axon overgrowth defect. Animals were considered as defective if DD6 axon terminal was located posteriorly to the rectum. *** p<0.001; n.s., not significant (Chi-square test). (**G**) Schematic of the relative positions of DD6, DA9, PDB neurons and *lin-44* expressing cells. Light graded orange represents hypothetical LIN-44 gradient. Scale bars: 10μm.

### *lin-44/wnt* instructs asymmetric localization of LIN-17/Fz and DSH-1/Dsh to the pruning neurites

Wnt ligands activate intracellular signaling cascade by inducing the clustering of the receptors. Consistently, Frizzled receptors are localized to the site of action in Wnt-dependent manners. For example, LIN-17/Fz is localized asymmetrically on the cell cortex during asymmetric cell division, to the posterior neurite of the PLM mechanosensory neuron during neuronal polarization, and to the posterior asynaptic axonal domain to inhibit presynaptic assembly (Goldstein et al., 2006; Hilliard and Bargmann, 2006; Klassen and Shen, 2007). We therefore examined the subcellular localization of the LIN-17::GFP fusion protein during PDB posterior neurite pruning. We observed significantly higher accumulation of LIN-17::GFP puncta at the tip of the posterior neurites compared with those at the anterior neurite at L2 stage (Figures 7A and 7D). The asymmetric LIN-17::GFP localization at the posterior neurites is dependent on *lin-44/wnt*: we did not observe significant LIN-17::GFP puncta in the *lin-44* null mutant animals (Figures 7B and 7D). In contrast, LIN-17::GFP puncta were accumulated at the posterior neurites of PDB in the *nrt-bfp-lin-44* animals (Figures 7C and 7D). This is consistent with no obvious pruning defects in the *nrt-bfp-lin-44* animals. We note that the LIN-17::GFP puncta observed in the ventral neurite of wildtype animals were completely absent in the *nrt-bfp-lin-44* animals (Figures 7A and 7C). These LIN-17::GFP puncta might be associated with the gradient-dependent function of *lin-44* such as neuronal polarization and guidance.

**Figure 7.**
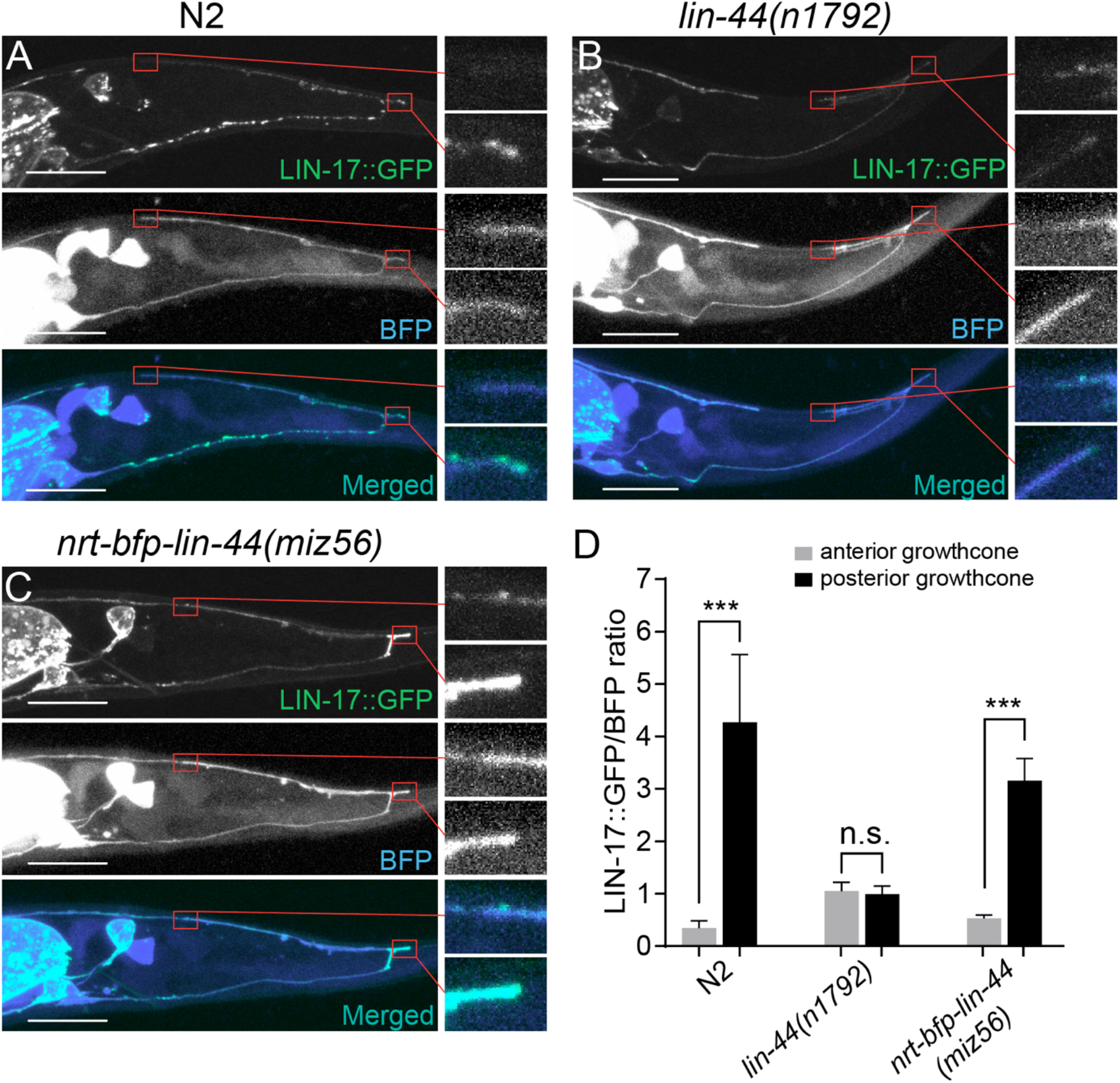
Wnt-dependent localization of LIN-17 at the PDB posterior neurites. (**A-C**) Representative images of LIN-17::GFP localization (top panels), PDB neurite structure labeled with cytoplasmic BFP (middle panels)and merged images (bottom panels) in N2 (**A**), *lin-44(n1792)* (**B**), and *nrt-bfp-lin-44(miz56)* (**C**) animals, respectively. Magnified images of the tip of anterior and posterior neurites are shown in the right panels. (**D**) Quantification of the normalized GFP/BFP signal ratio at the anterior and posterior growth cones. Error bars indicate mean ± SEM. ***p<0.001; n.s., not significant (Ratio paired t-test).

Dishevelled (Dsh/Dvl) proteins are the intracellular signaling components of both canonical and non-canonical Wnt signaling and are known to co-localize with active Frizzled receptors (Wallingford and Habas, 2005). There are three *Dishevelled* genes in *C.* elegans (*dsh-1, dsh-2* and *mig-5*). Similar to Frizzled receptors, Dsh proteins are often localized asymmetrically within the cells in Wnt-dependent manners (Heppert et al., 2018; Klassen and Shen, 2007; Mizumoto and Sawa, 2007; Walston et al., 2004). We did not observe significant pruning defects in *dsh-1* and *mig-5* single mutant animals, likely due to their functional redundancy (data not shown). Nevertheless, DSH-1/Dsh::GFP puncta were localized in the posterior neurites in the wildtype and *nrt-bfp-lin-44* but not in the *lin-44* null animals (Supplemental Figures 3A, 3B, 3D and 3E). Consistent with the idea that Dsh proteins are recruited to the active Frizzled receptors, DSH-1 localization at the posterior neurites was dependent on the LIN-17/Fz receptor (Supplemental Figures 3C and 3E). Taken these results together, we concluded that the locally restricted Wnt instructs asymmetric neurite pruning via recruiting Frizzled receptor and Dishevelled proteins to the pruning neurites (Figure 8).

**Figure 8.**
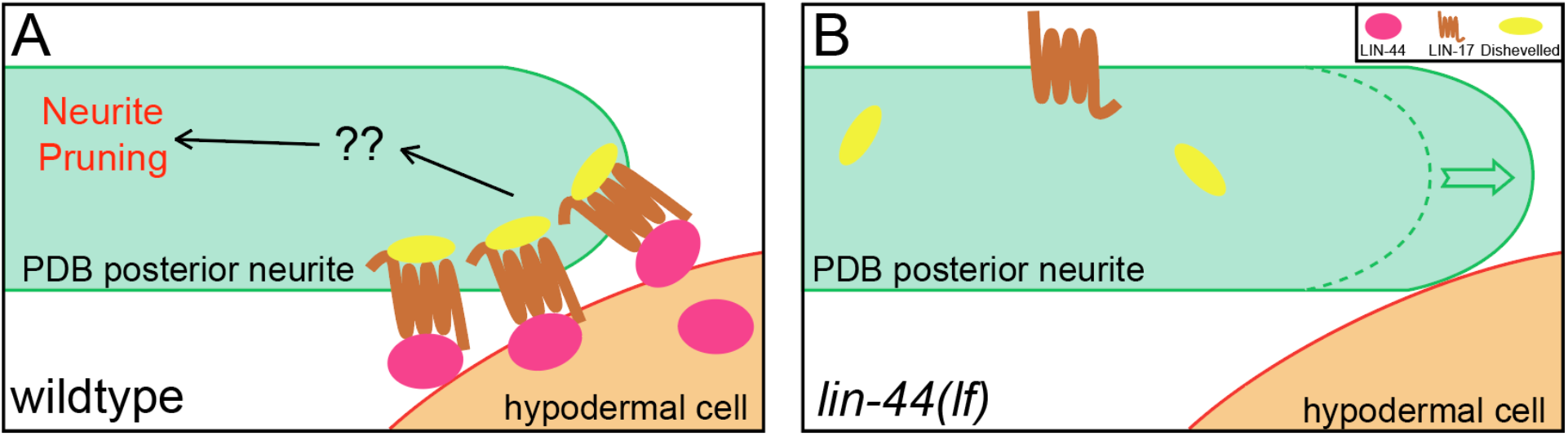
A model of asymmetric neurite pruning in PDB. LIN-44/Wnt from the tail hypodermal cell (hyp10) induces PDB neurite pruning via recruiting LIN-17/Fz and DSH-1/Dsh in wildtype (left panel) and defective pruning in *lin-44* mutants (right panel).

## Discussion

Developmental neurite pruning plays crucial roles in shaping functional neurocircuit, yet not much is known about the signaling cues that instruct stereotyped neurite pruning during neuronal development. Here, we found that Wnt signal from the hypodermal cells instructs stereotyped neurite pruning in the PDB cholinergic motorneuron in *C. elegans*. Asymmetric Wnt signal from the posterior hypodermal cells specifically induces pruning of the posterior neurite to sculpt the V-shape projection pattern of the PDB neurite. The unique structure of PDB and its topographic arrangement relative to the signaling cues allowed us to uncover the unique function of gradient-independent Wnt signaling in neurite pruning.

### Wnt signaling and neurite pruning

There have only been a handful of studies elucidating the roles of Wnt signaling in neurite pruning despite its critical roles in neuronal development and function. In *C. elegans*, two Wnts, CWN-1 and CWN-2, act through CAM-1/Ryk (receptor tyrosine kinase) to protect neurite from the *mbr-1-*mediated pruning in the AIM interneurons: the pruning defect in AIMs of *mbr-1* mutants is suppressed by mutations in *cam-1*, while overexpression of *cam-1* in AIMs inhibits neurite pruning (Hayashi et al., 2009). These data indicate that, in contrast to LIN-44/Wnt which instructs neurite pruning in PDB, CWN-1/2 inhibit neurite pruning in AIMs. It is possible that distinct receptors expressed in AIM and PDB neurons underlie the opposite effect of Wnt signaling on neurite pruning between them. Indeed, several studies have shown that *cam-1/Ror* and *lin-17/fz* function antagonistically (Goh et al., 2012; Kidd et al., 2015). The interesting analogy to our study has been observed in *Drosophila* contralaterally-projecting serotonin-immunoreactive deuterocerebral interneurons (CSDns) (Singh et al., 2010). CSDns undergo extensive dendritic remodeling during metamorphosis. The loss of Wingless and Wnt5 impairs the pruning of the transient dendritic branches of CSDn. Interestingly, the pruning of the CSDn dendrites is an activity-dependent process and requires their upstream neurons. Wnt signaling functions downstream of the neuronal activity to eliminate dendrites with no synaptic inputs from the upstream neurons. While it remains to be determined whether Wnt signaling plays an instructive or a permissive role in CSDn dendrite pruning, these observations along with our present study show conserved role of Wnt signaling as a pruning cue.

Wnt and Frizzled receptors control distinct downstream signaling cascades (Wnt-β-catenin; Wnt-PCP; Wnt-Ca^2+^) in context-dependent manners. We have examined potential downstream signaling components such as *bar-1/β-catenin*, *fmi-1/flamingo, vang-1/Van Gogh* but none of them showed significant neurite pruning defects (Figure 2E). Further candidate and forward genetic screenings will provide the mechanistic insights into the Wnt-dependent neurite pruning.

### Gradient-independent Wnt signaling

The normal development of *Drosophila* whose endogenous *wingless* is replaced with the membrane-tethered Wingless *(nrt-wg)* implies that gradient-independent or contact-dependent Wnt signaling is largely sufficient for the normal tissue patterning during development (Alexandre et al., 2014). The authors also noted the importance of the gradient signal of Wingless since the *nrt-wg* fly showed delayed development and reduced fitness. Consistently, recent works revealed the critical requirement of secretion and diffusion of Wingless for renal tube patterning and intestinal compartmentalization in *Drosophila* (Beaven and Denholm, 2018; Tian et al., 2019).

In this study, we showed that the membrane-tethered LIN-44 is sufficient for the PDB neurite pruning but not for DD6 axon termination and DA9 synapse patterning. How can cells distinguish gradient-dependent and independent Wnt signaling? One possible explanation is that each Wnt-dependent process requires a distinct threshold of the Wnt concentration to activate the signaling cascade. The processes that are gradient-dependent might require lower Wnt concentration to execute the process than the gradient-independent process. Alternatively, it is also possible that the gradient-independent or contact-dependent processes require transmembrane co-factors that mediate contact-dependent Wnt signaling. *sdn-1/syndecan* was one such candidate as it has been shown that *sdn-1* is required for the Wnt-dependent spindle orientation in early embryonic development (Dejima et al., 2014), and *sdn-1* mutants exhibit PDB structure defects similar to *lin-44/wnt* (Saied-Santiago et al., 2017). However, we did not observe significant pruning defects in PDB of *sdn-1* mutants (data not shown). Further candidate and forward genetic screenings will reveal novel factors that are specifically required for the contact-dependent Wnt signaling.

The membrane-tethered LIN-44/Wnt (Nrt-LIN-44) efficiently induced posterior neurite pruning, suggesting the contact with Wnt-expressing cells rather than the gradient distribution of LIN-44 triggers the PDB neurite pruning. The contact-dependent neurite pruning model can explain why the anterior neurite, which is also located in close proximity to the *lin-44-*expressing cells, can ignore the LIN-44/Wnt pruning cue. It also explains how the PDB neurite can initially grow posteriorly from its cell body toward the source of pruning cue (LIN-44-expressing cells). On the other hand, we do not exclude the possibility that the anterior neurite responds to the attractive guidance cues from the anterior cells so that it is protected from the Wnt-dependent neurite pruning. We have tested several candidates including *sax-7/Neurofascin*, which guides and stabilizes the dendritic arbor structure of the PVD sensory neuron (Dong et al., 2013), *unc-129/TGFβ* which is a neurite attractant expressed from the postsynaptic dorsal body wall muscles (Colavita, 1998), *unc-6/Netrin* that acts as an axon attractant through *unc-40/DCC* receptor (Chan et al., 1996), and *cam-1/Ryk* which acts downstream of Wnt to protect neurites from pruning (Hayashi et al., 2009), but none of them showed ectopic pruning of the anterior neurites. Future studies will be necessary to determine the threshold of Wnt concentration to induce neurite pruning as well as potential mechanisms that protect anterior neurite from pruning.

## Experimental procedures

### Strains

Bristol N2 strain was used as wildtype reference. All strains were cultured in the nematode growth medium (NGM) as described previously (Brenner, 1974) at 22°C. The following genotypes were used in this study: *lin-44(n1792), lin-44(miz56), lin-17(n671), egl-20(n585), mig-1(e1787), cam-1(gm122), bar-1(ga80), fmi-1(tm306), vang-1(ok1142), unc-13(e450), dsh-1(ok1445), mig-5(tm2639), sdn-1(ok449).* Genotyping primers are listed in the supplemental material.

### Transgenes

The transgenic lines were generated using standard microinjection method (Fire 1986; Mello et al. 1991): *mizIs9 (Pkal-1::zf1-GFPnovo2::CAAX; Pvha-6::zif-1; Pkal-1::mCherry::rab-3), wyIs486 (Pflp-13::2xGFP, Pplx-2::2xmCherry with Podr-1::RFP), mizEx248 (Plin-44::lin-44; Podr-1::RFP), mizEx349 (Plin-44::egl-20; Podr-1::RFP), mizEx247 (Pegl-20::lin-44, Podr-1::RFP), mizEx295 (Pkal-1::lin-17::GFPnovo2; Pkal-1::bfp; Podr-1::GFP), mizEx291 (Pkal-1::dsh-1::GFPnovo2; Pkal-1::bfp; Podr-1::GFP), mizEx317 (Plin-17::lin-17; Podr-1::RFP), mizEx374 (Prgef-1::lin-17; Podr-1::RFP), mizEx366 (Pmig-13::GFP::rab-3; Pmig-13::lin-17::mCherry; Podr-1::GFP), mizEx271 (Plin-44::BFP; Podr-1::GFP)*

### Plasmid construction

*C. elegans* expression clones were made in a derivative of pPD49.26 (A. Fire), the pSM vector (a kind gift from S. McCarroll and C. I. Bargmann). *dsh-1* cDNA clone was obtained by RT-PCR from N2 mRNA using Superscript III First-strand synthesis system and Phusion High-Fidelity DNA Polymerase (Thermo Fisher Scientific). *lin-17* cDNA clone was obtained from the plasmid used in the previous work (Mizumoto and Shen, 2013).

#### Pkal-1::zf1-GFPnovo2-caax plasmid

The 3.7kb fragment of *kal-1* promoter was amplified from N2 genomic DNA using Phusion high fidelity enzyme (ThermoFisher Scientific, USA) and cloned into the *Sph*I and *Asc*I sites of the pSM-GFPnovo2 vector (Hendi and Mizumoto, 2018). 111bp sequence of the ZF1 zinc finger domain from *pie-1* and 51bp sequence of the CAAX sequence from human KRas were inserted into the 5’ and 3’ of GFPnovo2 sequence, respectively, using Gibson assembly method (Armenti et al., 2014; Gibson et al., 2009). The intestinal GFP signal was reduced by expressing *zif-1* in the intestine under the *pha-6* promoter to degrade ZF1-GFPnovo2::CAAX.

#### Neurotactin-BFP-lin-44 repair plasmid

The plasmid containing a codon-optimized *neurotactin* cDNA with 2xHA tag was obtained from GeneArt (ThermoFisher Scientific, USA). *C. elegans* codon-optimized *BFP* with three synthetic introns was inserted into the 3’ end of the Neurotactin-2xHA sequence to generate *Neurotactin-2xHA-BFP* construct. 700bp of the 5’ homology arm of the *lin-44* promoter region, *Neurotactin-2xHA-BFP* (5’ portion until middle of intron 1), *BFP* (3’ portion from intron 1), 700bp of the 3’ homology arm spanning *lin-44* coding region are cloned into the *Sac*II and *Not*I sites of a dual-marker selection cassette (*loxP* + P*myo-2*::*GFP*::*unc-54* 3′UTR + P*rps-27*::*neoR*::*unc-54* 3′UTR *+ loxP* vector) (Au et al., 2019; Gibson et al., 2009).

#### *lin-44* gRNA constructs

Two 19bp gRNA sequences (gRNA3-CGATCAGTGGTGCACCTGC, gRNA4-TATTTCCGTCTTCAGCCAA) near the start codon of the *lin-44* gene were selected using CRISPR guide RNA selection tool (http://genome.sfu.ca/crispr/) and were cloned into *Rsa*I site of the sgRNA (F+E) vector, pTK73 (Obinata et al., 2018).

### CRISPR

The repair template plasmid, two *lin-44* sgRNA plasmids and a Cas9 plasmid (Addgene# 46168) (Friedland et al., 2013) were co-injected into young adults. The candidate genome-edited animals were screened based on G418 resistance and uniform expression of P*myo-2::GFP* in the pharynx as described previously (Au et al., 2019). The selection cassette was excised by injecting Cre recombinase plasmid (pDD104, Addgene #47551). Excision of the selection cassette, which was inserted within the first intron of BFP, reconstituted *Neurotactin-2xHA-BFP-lin-44.* The junctions between *neurotactin* and *BFP* as well as *BFP* and *lin-44* coding sequence were confirmed by Sanger sequencing.

### Semi-synchronization of *C. elegans*

Eggs at various stages were collected by bleaching the gravid adults with a bleaching solution (2.5% sodium hypochlorite and 0.5N Sodium Chloride) for 5 min. The eggs were washed twice with M9 buffer and seeded onto an NGM plate. For observing the pruning event, the animal was rescued from the slide after the first imaging at the L2 stage (approximately 26 hours after seeding eggs onto the NGM plates) and placed it in an individual NGM plate to resume the development for 24 hours before re-imaging at early L4 stages.

### Confocal microscopy

Images of fluorescently tagged fusion proteins were captured in live *C. elegans* using a Zeiss LSM800 Airyscan confocal microscope (Carl Zeiss, Germany). Worms were immobilized on 2% agarose pad using a mixture of 7.5 mM levamisole (Sigma-Aldrich) and 0.225M BDM (2,3-butanedione monoxime) (Sigma-Aldrich). Images were analyzed with Zen software (Carl Zeiss) or Image J (NIH, USA).

For time-lapse imaging, animals were mounted onto 5% agarose pads and immobilized using 0.5µl of 0.10 µmpolystyrene latex microsphere (Alfa Aesar # 427124Y). The coverslip was sealed with Vaseline (Vaseline® Jelly Original), to avoid dehydration of the animals and agarose pad during imaging.

### Statistics

Data were processed using Prism7 (GraphPad Software, USA). We applied the Chi-square test (with Yates’ continuity corrected) for comparison between two binary data groups, and one-way ANOVA method for comparison among more than three parallel groups with multiple plotting points. Data were plotted with error bars representing standard errors of the mean (SEM). *, ** and *** represent P-value <0.05, <0.01 and <0.001 respectively. Sample numbers were pre-determined before conducting statistical analyses.

## Acknowledgment

We are grateful to Don Moerman for suggestions and strains. We thank Harald Hutter, Kenji Sugioka and Riley St. Clair for critical reading of the manuscript, Ardalan Hendi for generating some strains, Shinsuke Niwa for the pTK73 plasmid, and the Mizumoto lab members for general discussions. Some strains used in this study are obtained from the CGC, which is funded by NIH Office of Research Infrastructure Programs (P40 OD010440), and from the National Bioresource Project, Japan. This project is funded by HFSP (CDA-00004/2014) and NSERC (RGPIN-2015-04022). KM is a Tier 2 Canada Research Chair and a Michael Smith Foundation for Health Research scholar.

## Figures

**Supplemental Figure 1.**
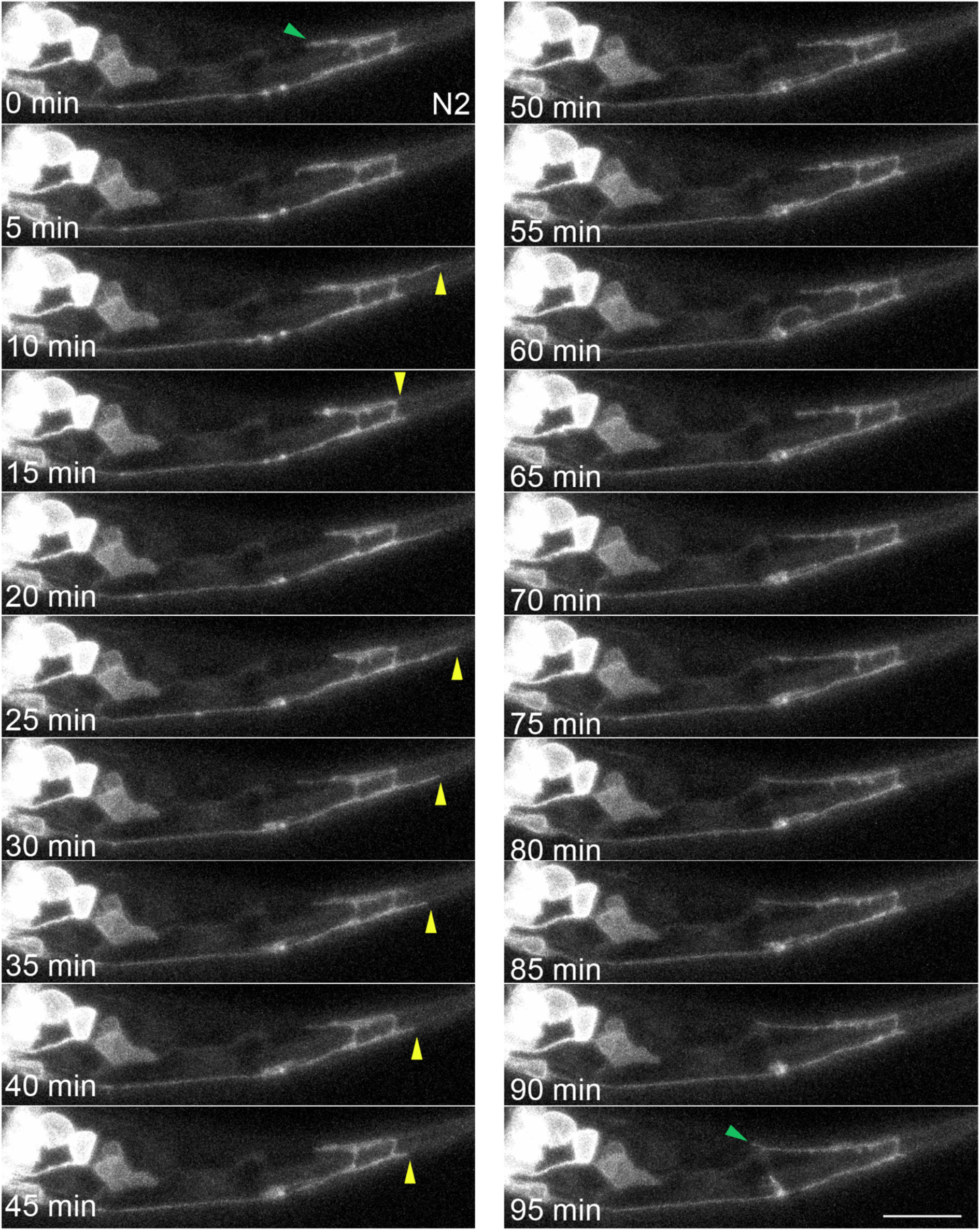
Time-lapse imaging of PDB neurite pruning. PDB neurite is labeled with *mizIs9* marker. Images were taken every 5 minutes. Green arrows denote the end of anterior neurite in the first and last frame, and yellow arrows denote the end of posterior neurites during two pruning events. Scale bars: 10 μm.

**Supplemental Figure 2.**
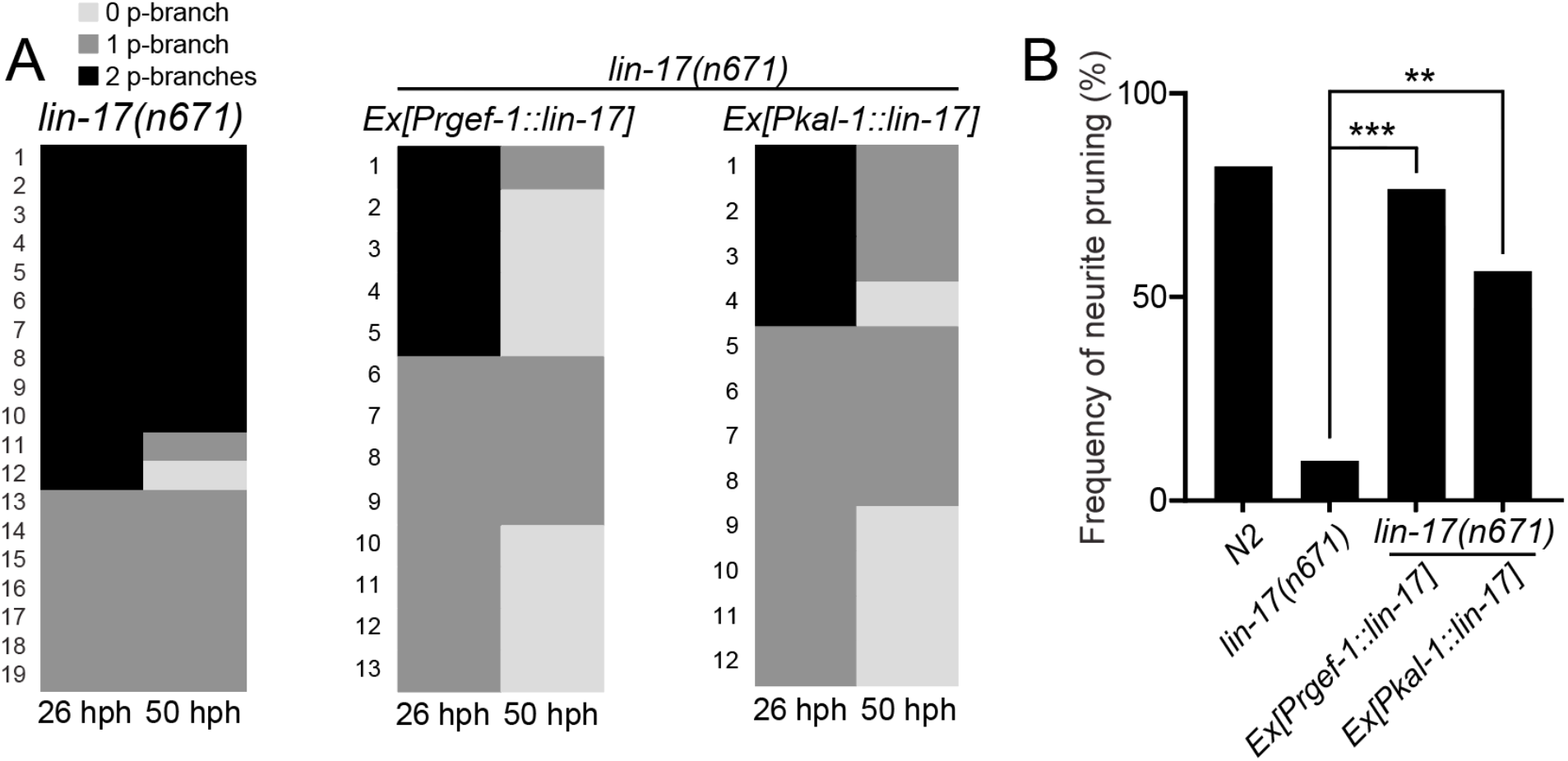
*lin-17/fz* acts cell-autonomously in PDB. (**A**) Quantification of the posterior neurite numbers of individual animals at L2 and L4 stages in *lin-17* mutants and *lin-17* mutants with rescuing transgenes. Note that quantification of *lin-17* mutants is from Figure 3. (**B**) Quantification of posterior neurite pruning frequency. *** p < 0.001; ** p < 0.002 (Chi-square test with Yates’ correction).

**Supplemental Figure 3.**
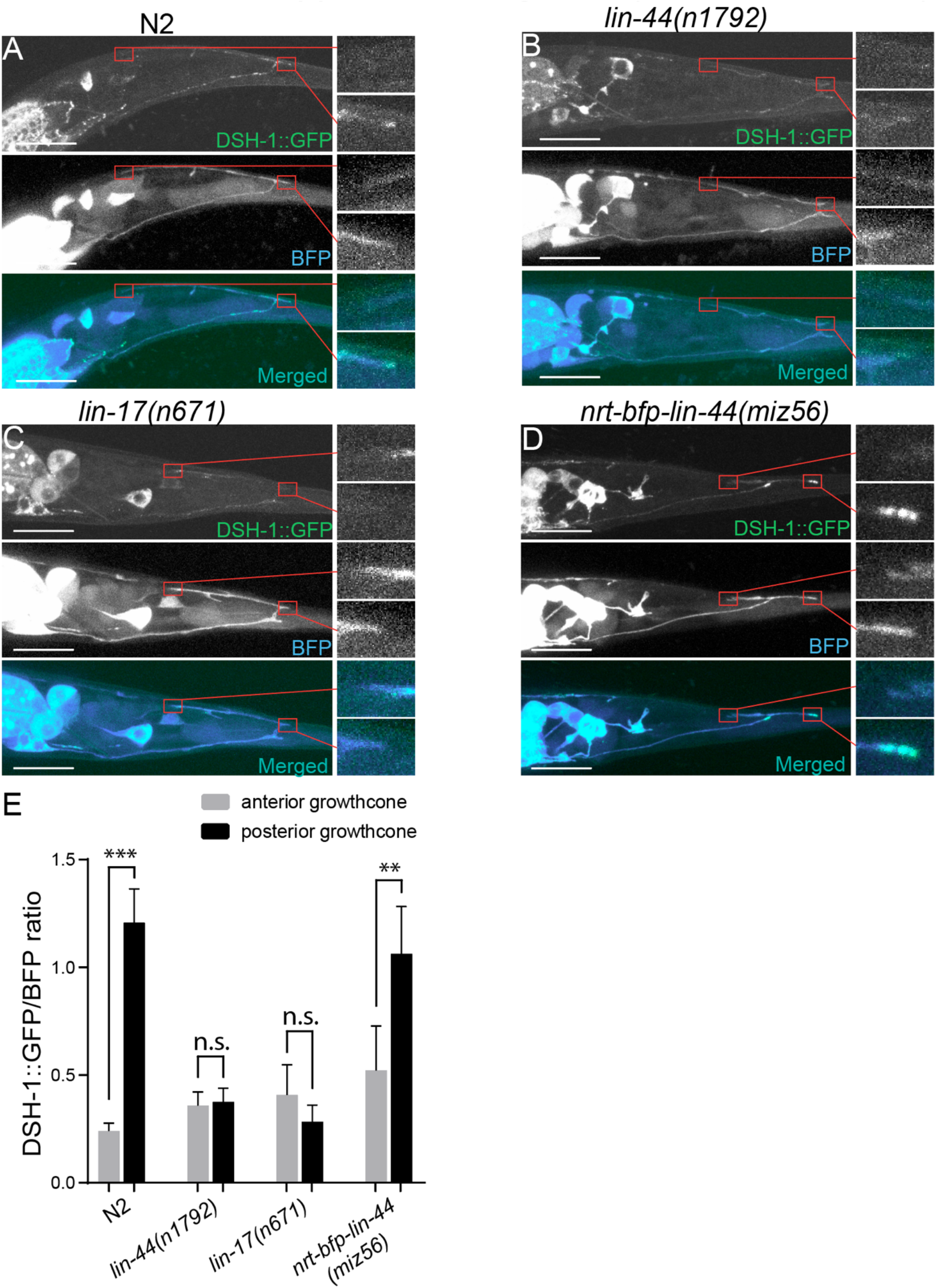
Wnt-dependent localization of DSH-1/Dsh at the PDB posterior neurites. (**A-D**) Representative images of DSH-1::GFP localization (top panels), PDB neurite structure labeled with cytoplasmic BFP (middle panels) and merged images (bottom panels) in N2 (**A**), *lin-44(n1792)* (**B**), *lin-17(n671)* (**C**) and *nrt-bfp-lin-44(miz56)* (**D**) animals, respectively. Magnified images of the tip of anterior and posterior neurites are shown in the right panels. (**E**) Quantification of the normalized GFP/BFP signal ratio at the anterior and posterior growth cones. Error bars indicate mean ± SEM. ***p<0.001; n.s., not significant (Ratio paired t-test).

## Supplemental Information

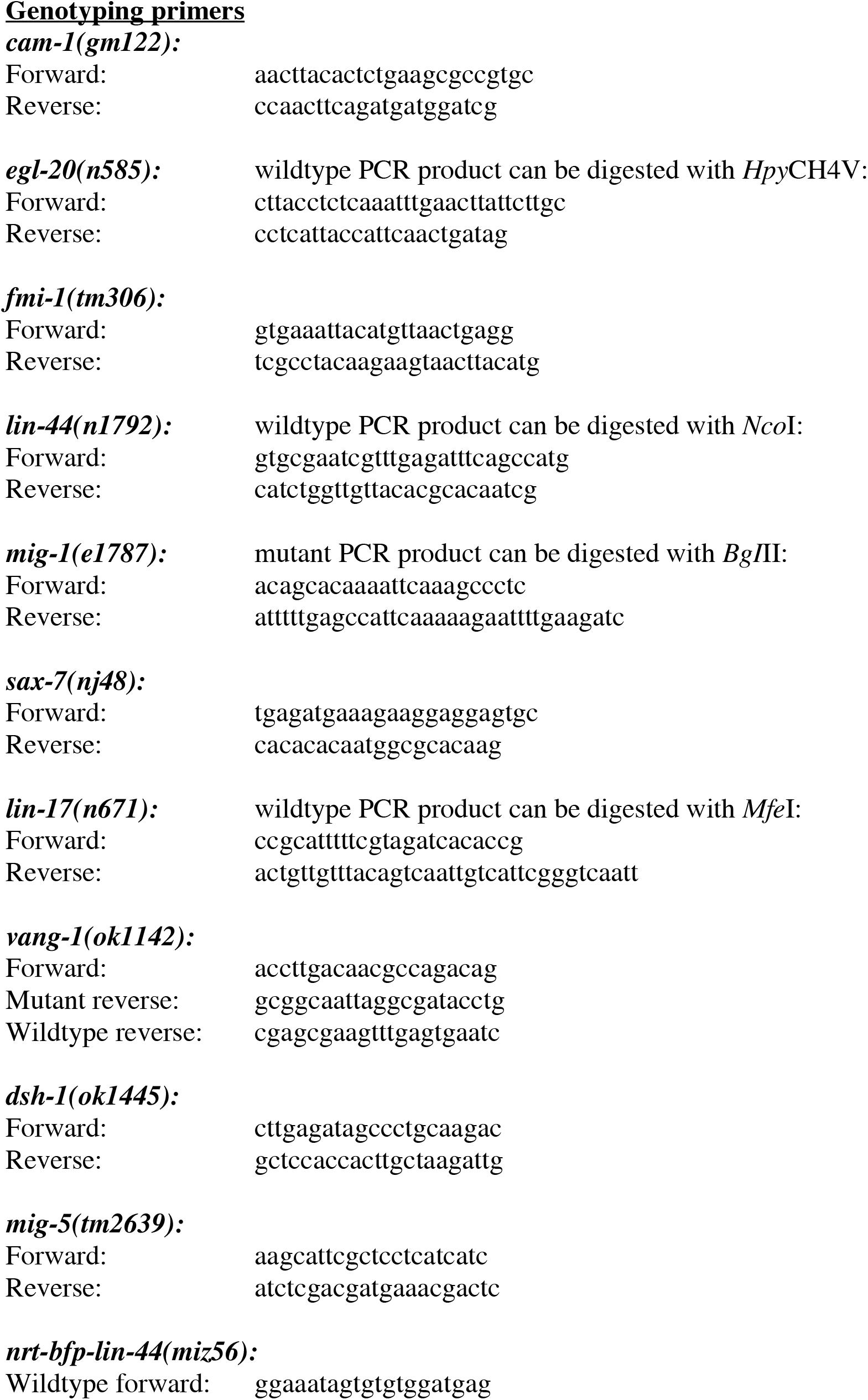
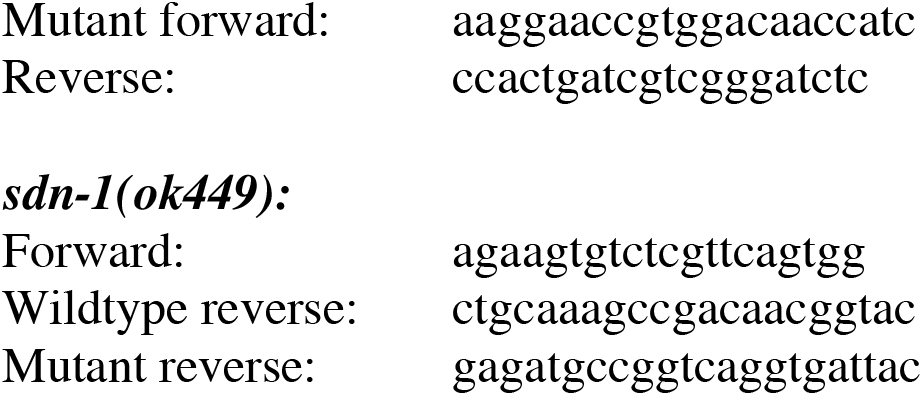

## References

Alexandre C, Baena-Lopez A, Vincent JP. 2014. Patterning and growth control by membrane-tethered wingless. Nature 505:180–185. doi:10.1038/nature12879

Armenti ST, Lohmer LL, Sherwood DR, Nance J. 2014. Repurposing an endogenous degradation system for rapid and targeted depletion of C. elegans proteins. Development 141:4640–4647. doi:10.1242/dev.115048

Au V, Li-Leger E, Raymant G, Flibotte S, Chen G, Martin K, Fernando L, Doell C, Rosell FI, Wang S, Edgley ML, Rougvie AE, Hutter H, Moerman DG. 2019. CRISPR/Cas9 Methodology for the Generation of Knockout Deletions in Caenorhabditis elegans. G3 Genes|Genomes|Genetics 9:135–144. doi:10.1534/g3.118.200778

Baena-Lopez LA, Franch-Marro X, Vincent J-P. 2009. Wingless Promotes Proliferative Growth in a Gradient-Independent Manner. Sci Signal 2:ra60–ra60. doi:10.1126/scisignal.2000360

Bagri A, Cheng HJ, Yaron A, Pleasure SJ, Tessier-Lavigne M. 2003. Stereotyped pruning of long hippocampal axon branches triggered by retraction inducers of the semaphorin family. Cell 113:285–299. doi:10.1016/S0092-8674(03)00267-8

Bartscherer K, Boutros M. 2008. Regulation of Wnt protein secretion and its role in gradient formation. EMBO Rep 9:977–982. doi:10.1038/embor.2008.167

Beaven R, Denholm B. 2018. Release and spread of Wingless is required to pattern the proximo-distal axis of Drosophila renal tubules. Elife 7:1–17. doi:10.7554/eLife.35373

Bishop DL, Misgeld T, Walsh MK, Gan W-B, Lichtman JW. 2004. Axon Branch Removal at Developing Synapses by Axosome Shedding. Neuron 44:651–661. doi:10.1016/j.neuron.2004.10.026

Brenner S. 1974. The genetics of Caenorhabditis elegans. Genetics 77:71–94. doi:10.1002/cbic.200300625

Bülow HE, Berry KL, Topper LH, Peles E, Hobert O. 2002. Heparan sulfate proteoglycan-dependent induction of axon branching and axon misrouting by the Kallmann syndrome gene kal-1. Proc Natl Acad Sci 99:6346–6351. doi:10.1073/pnas.092128099

Chan SS-Y, Zheng H, Su M-W, Wilk R, Killeen MT, Hedgecock EM, Culotti JG. 1996. UNC-40, a C. elegans Homolog of DCC (Deleted in Colorectal Cancer), Is Required in Motile Cells Responding to UNC-6 Netrin Cues. Cell 87:187–195. doi:10.1016/S0092-8674(00)81337-9

Colavita A. 1998. Pioneer Axon Guidance by UNC-129, a C. elegans TGF-. Science (80-) 281:706–709. doi:10.1126/science.281.5377.706

Colman H, Nabekura J, Lichtman JW. 1997. Alterations in synaptic strength preceding axon withdrawal. Science 275:356–61. doi:10.1126/science.275.5298.356

Coudreuse DYM. 2006. Wnt Gradient Formation Requires Retromer Function in Wnt-Producing Cells. Science (80-) 312:921–924. doi:10.1126/science.1124856

Dejima K, Kang S, Mitani S, Cosman PC, Chisholm AD. 2014. Syndecan defines precise spindle orientation by modulating Wnt signaling in C. elegans. J Cell Sci 127:e1–e1. doi:10.1242/jcs.165761

Deppmann CD, Mihalas S, Sharma N, Lonze BE, Niebur E, Ginty DD. 2008. A Model for Neuronal Competition During Development. Science (80-) 320:369–373. doi:10.1126/science.1152677

Dong X, Liu OW, Howell AS, Shen K. 2013. An Extracellular Adhesion Molecule Complex Patterns Dendritic Branching and Morphogenesis. Cell 155:296–307. doi:10.1016/j.cell.2013.08.059

Eisenmann DM. 2005. Wnt signaling. WormBook. doi:10.1895/wormbook.1.7.1

Friedland AE, Tzur YB, Esvelt KM, Colaiácovo MP, Church GM, Calarco JA. 2013. Heritable genome editing in C. elegans via a CRISPR-Cas9 system. Nat Methods 10:741–3. doi:10.1038/nmeth.2532

Gibson DG, Young L, Chuang R-Y, Venter JC, Hutchison CA, Smith HO. 2009. Enzymatic assembly of DNA molecules up to several hundred kilobases. Nat Methods 6:343–345. doi:10.1038/nmeth.1318

Goh KY, Ng NW, Hagen T, Inoue T. 2012. p21-Activated kinase interacts with Wnt signaling to regulate tissue polarity and gene expression. Proc Natl Acad Sci 109:15853–15858. doi:10.1073/pnas.1120795109

Goldstein B, Takeshita H, Mizumoto K, Sawa H. 2006. Wnt signals can function as positional cues in establishing cell polarity. Dev Cell 10:391–396. doi:10.1016/j.devcel.2005.12.016

Gross JC, Chaudhary V, Bartscherer K, Boutros M. 2012. Active Wnt proteins are secreted on exosomes. Nat Cell Biol 14:1036–1045. doi:10.1038/ncb2574

Hayashi Y, Hirotsu T, Iwata R, Kage-Nakadai E, Kunitomo H, Ishihara T, Iino Y, Kubo T. 2009. A trophic role for Wnt-Ror kinase signaling during developmental pruning in Caenorhabditis elegans. Nat Neurosci 12:981–7. doi:10.1038/nn.2347

He C, Liao C, Pan C. 2018. Wnt signalling in the development of axon, dendrites and synapses. Open Biol 8:180116. doi:10.1098/rsob.180116

Hendi A, Mizumoto K. 2018. GFPnovo2, a brighter GFP variant for in vivo labeling in C. elegans. microPublication Biol. 10.17912/49YB-7K39

Heppert JK, Pani AM, Roberts AM, Dickinson DJ, Goldstein B. 2018. A CRISPR Tagging-Based Screen Reveals Localized Players in Wnt-Directed Asymmetric Cell Division. Genetics 208:1147–1164. doi:10.1534/genetics.117.300487

Herman MA, Vassilieva LL, Horvitz HR, Shaw JE, Herman RK. 1995. The C. elegans gene lin-44, which controls the polarity of certain asymmetric cell divisions, encodes a Wnt protein and acts cell nonautonomously. Cell 83:101–110. doi:10.1016/0092-8674(95)90238-4

Hilliard MA, Bargmann CI. 2006. Wnt signals and Frizzled activity orient anterior-posterior axon outgrowth in C. elegans. Dev Cell 10:379–390. doi:10.1016/j.devcel.2006.01.013

Inestrosa NC, Varela-Nallar L. 2015. Wnt signalling in neuronal differentiation and development. Cell Tissue Res 359:215–223. doi:10.1007/s00441-014-1996-4

Jan Y-N, Jan LY. 2010. Branching out: mechanisms of dendritic arborization. Nat Rev Neurosci 11:316–328. doi:10.1038/nrn2836

Kage E, Hayashi Y, Takeuchi H, Hirotsu T, Kunitomo H, Inoue T, Arai H, Iino Y, Kubo T. 2005. MBR-1, a novel helix-turn-helix transcription factor, is required for pruning excessive neurites in Caenorhabditis elegans. Curr Biol 15:1554–1559. doi:10.1016/j.cub.2005.07.057

Kidd AR, Muñiz-Medina V, Der CJ, Cox AD, Reiner DJ. 2015. The C. Elegans Chp/Wrch ortholog CHW-1 contributes to LIN-18/Ryk and LIN-17/ frizzled signaling in cell polarity. PLoS One 10:1–21. doi:10.1371/journal.pone.0133226

Klassen MP, Shen K. 2007. Wnt Signaling Positions Neuromuscular Connectivity by Inhibiting Synapse Formation in C. elegans. Cell 130:704–716. doi:10.1016/j.cell.2007.06.046

Liu Y, Rutlin M, Huang S, Barrick CA, Wang F, Jones KR, Tessarollo L, Ginty DD. 2012. Sexually Dimorphic BDNF Signaling Directs Sensory Innervation of the Mammary Gland. Science (80-) 338:1357–1360. doi:10.1126/science.1228258

Maro GS, Klassen MP, Shen K. 2009. A β-Catenin-Dependent Wnt Pathway Mediates Anteroposterior Axon Guidance in C. elegans Motor Neurons. PLoS One 4:e4690. doi:10.1371/journal.pone.0004690

Mizumoto K, Sawa H. 2007. Cortical β-Catenin and APC Regulate Asymmetric Nuclear β-Catenin Localization during Asymmetric Cell Division in C. elegans. Dev Cell 12:287–299. doi:10.1016/j.devcel.2007.01.004

Mizumoto K, Shen K. 2013. Two Wnts Instruct Topographic Synaptic Innervation in C. elegans. Cell Rep 5:389–396. doi:10.1016/j.celrep.2013.09.011

Nusse R. 2005. Wnt signaling in disease and in development. Cell Res 15:28–32. doi:10.1038/sj.cr.7290260

Obinata H, Sugimoto A, Niwa S. 2018. Streptothricin acetyl transferase 2 (Sat2): A dominant selection marker for Caenorhabditis elegans genome editing. PLoS One 13:e0197128. doi:10.1371/journal.pone.0197128

Pani AM, Goldstein B. 2018. Direct visualization of a native Wnt in vivo reveals that a long-range Wnt gradient forms by extracellular dispersal. Elife 7:1–22. doi:10.7554/elife.38325

Park M, Shen K. 2012. WNTs in synapse formation and neuronal circuitry. EMBO J 31:2697– 2704. doi:10.1038/emboj.2012.145

Riccomagno MM, Hurtado A, Wang H, Macopson JGJ, Griner EM, Betz A, Brose N, Kazanietz MG, Kolodkin AL. 2012. The RacGAP β2-chimaerin selectively mediates axonal pruning in the hippocampus. Cell 149:1594–1606. doi:10.1016/j.cell.2012.05.018

Riccomagno MM, Kolodkin AL. 2015. Sculpting Neural Circuits by Axon and Dendrite Pruning. Annu Rev Cell Dev Biol 31:779–805. doi:10.1146/annurev-cellbio-100913-013038

Richmond JE, Davis WS, Jorgensen EM. 1999. UNC-13 is required for synaptic vesicle fusion in C. elegans. Nat Neurosci 2:959–964. doi:10.1038/14755

Routledge D, Scholpp S. 2019. Mechanisms of intercellular Wnt transport. Development 146:dev176073. doi:10.1242/dev.176073

Saha S, Aranda E, Hayakawa Y, Bhanja P, Atay S, Brodin NP, Li J, Asfaha S, Liu L, Tailor Y, Zhang J, Godwin AK, Tome WA, Wang TC, Guha C, Pollard JW. 2016. Macrophage-derived extracellular vesicle-packaged WNTs rescue intestinal stem cells and enhance survival after radiation injury. Nat Commun 7:13096. doi:10.1038/ncomms13096

Sahay A, Molliver ME, Ginty DD, Kolodkin AL. 2003. Semaphorin 3F Is Critical for Development of Limbic System Circuitry and Is Required in Neurons for Selective CNS Axon Guidance Events. J Neurosci 23:6671–6680. doi:10.1523/JNEUROSCI.23-17-06671.2003

Saied-Santiago K, Townley RA, Attonito JD, da Cunha DS, Díaz-Balzac CA, Tecle E, Bülow HE. 2017. Coordination of Heparan Sulfate Proteoglycans with Wnt Signaling To Control Cellular Migrations and Positioning in Caenorhabditis elegans. Genetics 206:1951–1967. doi:10.1534/genetics.116.198739

Sar Shalom H, Goldner R, Golan-Vaishenker Y, Yaron A. 2019. Balance between BDNF and Semaphorins gates the innervation of the mammary gland. Elife 8:1–20. doi:10.7554/eLife.41162

Sawa H, Korswagen HC. 2013. Wnt signaling in C. elegans. WormBook 1–30. doi:10.1895/wormbook.1.7.2

Sawa H, Lobel L, Horvitz HR. 1996. The Caenorhabditis elegans gene lin-17, which is required for certain asymmetric cell divisions, encodes a putative seven-transmembrane protein similar to the Drosophila Frizzled protein. Genes Dev 10:2189–2197. doi:10.1101/gad.10.17.2189

Schuldiner O, Yaron A. 2015. Mechanisms of developmental neurite pruning. Cell Mol Life Sci 72:101–119. doi:10.1007/s00018-014-1729-6

Singh AP, VijayRaghavan K, Rodrigues V. 2010. Dendritic refinement of an identified neuron in the Drosophila CNS is regulated by neuronal activity and Wnt signaling. Development 137:1351–1360. doi:10.1242/dev.044131

Stanganello E, Hagemann AIH, Mattes B, Sinner C, Meyen D, Weber S, Schug A, Raz E, Scholpp S. 2015. Filopodia-based Wnt transport during vertebrate tissue patterning. Nat Commun 6:1–14. doi:10.1038/ncomms6846

Sulston JE, Horvitz HR. 1977. Post-embryonic cell lineages of the nematode, Caenorhabditis elegans. Dev Biol 56:110–156. doi:10.1016/0012-1606(77)90158-0

Tian A, Duwadi D, Benchabane H, Ahmed Y. 2019. Essential long-range action of Wingless/Wnt in adult intestinal compartmentalization. PLOS Genet 15:e1008111. doi:10.1371/journal.pgen.1008111

van Amerongen R, Nusse R. 2009. Towards an integrated view of Wnt signaling in development. Development 136:3205–3214. doi:10.1242/dev.033910

Walker DS, Yan G, Towlson EK, Schafer WR, Barabási A-L, Chew YL, Vértes PE. 2017. Network control principles predict neuron function in the Caenorhabditis elegans connectome. Nature 550:519–523. doi:10.1038/nature24056

Wallingford JB, Habas R. 2005. The developmental biology of Dishevelled: an enigmatic protein governingcell fate and cell polarity. Development 132:4421–4436. doi:10.1242/dev.02068

Walston T, Tuskey C, Edgar L, Hawkins N, Ellis G, Bowerman B, Wood W, Hardin J. 2004. Multiple Wnt signaling pathways converge to orient the mitotic spindle in early C. elegans embryos. Dev Cell 7:831–841. doi:10.1016/j.devcel.2004.10.008

Whangbo J, Kenyon C. 1999. A Wnt Signaling System that Specifies Two Patterns of Cell Migration in C. elegans. Mol Cell 4:851–858. doi:10.1016/S1097-2765(00)80394-9

Yamamoto Y, Takeshita H, Sawa H. 2011. Multiple Wnts Redundantly Control Polarity Orientation in Caenorhabditis elegans Epithelial Stem Cells. PLoS Genet 7:e1002308. doi:10.1371/journal.pgen.1002308

Yang Y. 2012. Wnt signaling in development and disease. Cell Biosci 2:14. doi:10.1186/2045-3701-2-14

Zecca M, Basler K, Struhl G. 1996. Direct and Long-Range Action of a Wingless Morphogen Gradient. Cell 87:833–844. doi:10.1016/S0092-8674(00)81991-1

